# Hhex regulates the specification and growth of the hepatopancreatic ductal system

**DOI:** 10.1101/330779

**Authors:** Alethia Villasenor, Sébastien Gauvrit, Michelle M. Collins, Silvia Parajes, Hans-Martin Maischein, Didier Stainier

## Abstract

Significant efforts have advanced our understanding of foregut-derived organ development; however, little is known about the molecular mechanisms that underlie the formation of the hepatopancreatic ductal (HPD) system. Here, we report a role for the homeodomain transcription factor Hhex in directing HPD progenitor specification in zebrafish. Loss of Hhex function results in impaired HPD system formation. We found that Hhex specifies a distinct population of HPD progenitors that gives rise to the cystic duct, common bile duct, and extra-pancreatic duct. Since *hhex* is not uniquely expressed in the HPD region but is also expressed in endothelial cells and the yolk syncytial layer (YSL), we tested the role of blood vessels as well as the YSL in HPD formation. We found that blood vessels are required for HPD patterning, but not for HPD progenitor specification. In addition, we found that Hhex is required in both the endoderm and the YSL for HPD development. Our results shed light on the mechanisms necessary to direct endodermal progenitors towards the HPD fate and also advance our understanding of HPD system formation.

## INTRODUCTION

Cells that reside in the hepatopancreatic ductal (HPD) system have recently come under renewed interest because of their ability to differentiate into other endodermal cell lineages [1, 2]. Isolated HPD cells share the characteristics of stem cells as they can remain undifferentiated for months in culture [3], and can also be differentiated into hepatocytes, cholangiocytes or pancreatic endocrine cells depending on culture conditions [4, 5]. In addition, under pancreatic regenerative conditions, committed endocrine progenitors have been identified in glands of the HPD system indicating that HPD cells can differentiate into endocrine cells [1, 6]. These endocrine cells aggregate and form islet-like clusters in the liver parenchyma and can express mature endocrine markers including insulin, somatostatin, pancreatic polypeptide, and glucagon. It has thus been proposed that the HPD system acts as a reservoir for multipotent cells [4]. Elucidating the mechanisms responsible for HPD specification is likely to help advance reprogramming studies and regenerative medicine therapies.

The HPD system is composed of the extrapancreatic duct (EPD), common bile duct (CBD), cystic duct (CD) and extrahepatic duct (EHD) [7, 8]. It serves to connect the liver, gallbladder, and pancreas to the intestinal tract as well as to transport fluids between these organs for the adequate function of the digestive system. The mechanisms that direct HPD fate and the formation of its different components are only partially understood, with only a few signaling molecules known to be implicated in the process [9]. Among them, *Sox17* is known to play an important role in murine HPD development [10, 11]. *Sox17* segregates ductal progenitors from pancreatic progenitors in the ventral foregut thereby acting as a switch between the biliary and pancreatic fates. Misexpression of *Sox17* is sufficient to induce ductal fate while loss of *Sox17* leads to ectopic expression of PDX1^+^ cells in the liver and CBD [11]. However, loss of *Sox17* only leads to the absence of the gallbladder and CD suggesting that multiple transcription factors regulate the specification of the various components of the HPD system.

Another transcription factor that has been implicated in biliary development is Hhex [12-15]. In mouse, HHEX is expressed in ductal cells of the HPD system and pancreas [14] and absence of *Hhex* function causes a lack of pseudostratification of the hepatic diverticulum, defects in hepatoblast migration and impaired liver development [12, 13]. In addition, specific deletion of *Hhex* from the hepatic diverticulum leads to defects in hepatic biliary development, absence of gallbladder formation, and irregular morphogenesis of the extrahepatic biliary tract [14]. Here, we further investigate the role of *Hhex* in HPD development using the zebrafish model. The zebrafish model offers many advantages for the analysis of HPD development including its ex-utero development, optical transparency, as well as the ability of embryos to survive without a functional cardiovascular or digestive system [16, 17]. We therefore generated a *hhex* mutant line and found that *hhex* mutants do not develop an HPD system. Our results show that Hhex function is necessary for the specification of a pool of HPD progenitors and the subsequent formation of the CD, CBD, and EPD and that both cell-autonomous and cell non-autonomous functions of Hhex are necessary for HPD development. Our studies also identify a previously unreported role for the vasculature in modulating the patterning of the HPD system.

## RESULTS

### Generation of *hhex* mutants

To study the role of *hhex* in HPD system formation, we generated two *hhex* mutant alleles, *s721* and *s722*, using transcription activator-like effector nucleases (TALENs) [18-20]. The TALEN pairs were directed towards exon 3, and both mutant alleles contain lesions in the DNA binding domain of Hhex (**Fig. S1A**). The *s722* allele is an in-frame deletion which removes three amino acids (R149 to A151) from the homeodomain, while the *s721* allele consists of a 10 base pair insertion that leads to a premature stop codon (p.R149fs*2) and is predicted to remove 46% of the homeodomain as well as the complete C-terminal domain of Hhex (**Fig. S1B**). Homozygous carriers of either mutant allele do not survive beyond the larval stage and die starting at 8 days post fertilization (dpf).

We first investigated the liver and pancreas in *hhex*^*s721*^ mutants, hereafter abbreviated as *hhex*^*-/-*^ mutants, as defects in both of these organs have previously been reported in mouse, *Xenopus*, and zebrafish [12, 14, 21-25]. Similar to the results reported before for loss of HHEX function in mouse, *hhex* mutants display impaired liver growth and lack of the exocrine pancreas [12, 25]. Incrosses of *Tg(fabp:GFP); hhex*^*+/-*^ and *Tg(prox1a*:*RFP*); *hhex*^+/-^ show that mutant larvae have smaller and under-developed livers (**Fig. 1A-B**). In addition, whole-mount *in situ* hybridizations for *ptf1a* expression to mark the exocrine pancreas confirmed a complete absence of this tissue in *hhex* mutants (**Fig. 1C**). These endodermal phenotypes are present in both alleles (data not shown) and show full penetrance and expressivity.

**Figure 1.**
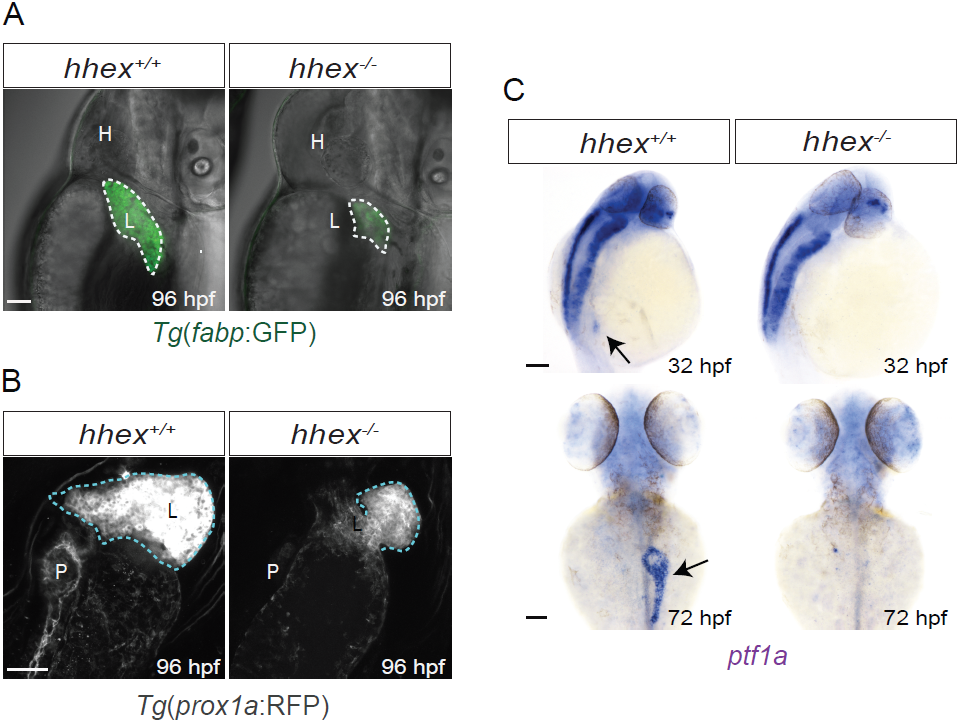
*hhex* mutants display impaired liver and pancreas development. **(A-C)** Analysis of the endodermal phenotypes in *hhex* mutants. **(A)** Lateral views of 96 hpf wild-type and *hhex* mutant larvae.**(B)** Maximum intensity projections of 96 hpf wild-type and *hhex* mutant. Ventral views, anterior to the top. *hhex* mutants display impaired liver development. **(C)** Dorsal views of whole-mount *in situ* hybridization for *ptf1a* expression at 32 and 72 hpf. Compared to stage-matched wild-type, *hhex* mutants lack exocrine tissue specification, but they maintain *ptf1a* neural expression. Black arrows point to pancreatic exocrine tissue. L, liver; P, pancreas; H, heart. Scale bars: 50 μm.

It has recently been reported that *Hhex* is necessary to maintain the differentiated state of δ-cells (i.e., Somatostatin (Sst) producing cells) in mouse [22]. *Hhex*-deficient islets contained fewer Sst^+^ cells and decreased Sst secretion. In addition, deletion of *Hhex* specifically within endocrine progenitors led to a complete loss of δ-cells in embryonic and postnatal mouse islets [22]. In order to check whether this function of Hhex was conserved in zebrafish, we analyzed incrosses of *Tg(sst:RFP); hhex*^*+/-*^ larvae at 80 hpf (**Fig. S2A-B**). We found that Hhex-deficient larvae had fewer *sst*:RFP^+^ cells (21± 3.6 in control n=12 embryos total, composed of wild-type n=4 and *hhex*^*+/-*^ n=8, compared to 14± 3.9 in *hhex* mutants, n=10 embryos; p=0.000174) and that these cells exhibited a dimmer RFP signal than wild-type cells. We also performed Sst immunostaining in 80 hpf larvae from *hhex*^+/-^ incrosses and observed significantly fewer Sst^+^ cells (21± 1.8 in control siblings n=5 compared to 1.8 ± 0.85 in *hhex* mutants, n=4; p=5E-05) (**Fig. S2C-D**). Interestingly, *hhex* mutants exhibited an increase in Insulin^+^ cells (β-cells) (34 ± 8.3 n=12 embryos total, composed of wild-type n=4 and *hhex*^*+/-*^ n=8, compared to 45 ± 11.9 in *hhex* mutants, n=10 embryos; p=0.0238) and Glucagon^+^ (α-cells) (20 ± 3.9 in control siblings n=12 embryos total, composed of wild-type n=4 and *hhex*^*+/-*^ n=8, compared to 40 ± 8.0 in *hhex* mutants, n=10 embryos; p=1.68E-07) (**Fig. S2B**). Our results show that Hhex is necessary to maintain fully differentiated δ-cells in zebrafish.

### *hhex* is expressed in the HPD system and regulates its morphogenesis

To study the role of *hhex* in the ontogeny of the HPD system, we first analyzed the spatiotemporal pattern of *hhex* expression in the HPD and surrounding tissues. Whole-mount *in situ* hybridization revealed that before 32 hours post-fertilization (hpf), *hhex* is strongly expressed in blood endothelial cells. Its expression in the endoderm starts at around 20 hpf when the first *hhex*^+^ cells appear at the midline populating the nascent dorsal pancreatic area. By 24 hpf, *hhex*^+^ cells are also found slightly anterior to the dorsal pancreas in the liver region, and by 32 hpf *hhex* is expressed in the HPD progenitors (**Fig. 2A**, red arrow). Notably at this stage, *hhex* expression in blood endothelial cells becomes significantly reduced, while expression in the endoderm strongly increases. By 50 hpf, *hhex* expression expands towards the HPD primordium (**Fig. 2A**, blue arrow) and by 72 hpf *hhex* is expressed in all components of the HPD system: the EHD, CD, CBD, and EPD, as well as in the gallbladder, major intrahepatic ducts (IHD), and some cells of the principal islet (**Fig. 2A**).

**Figure 2.**
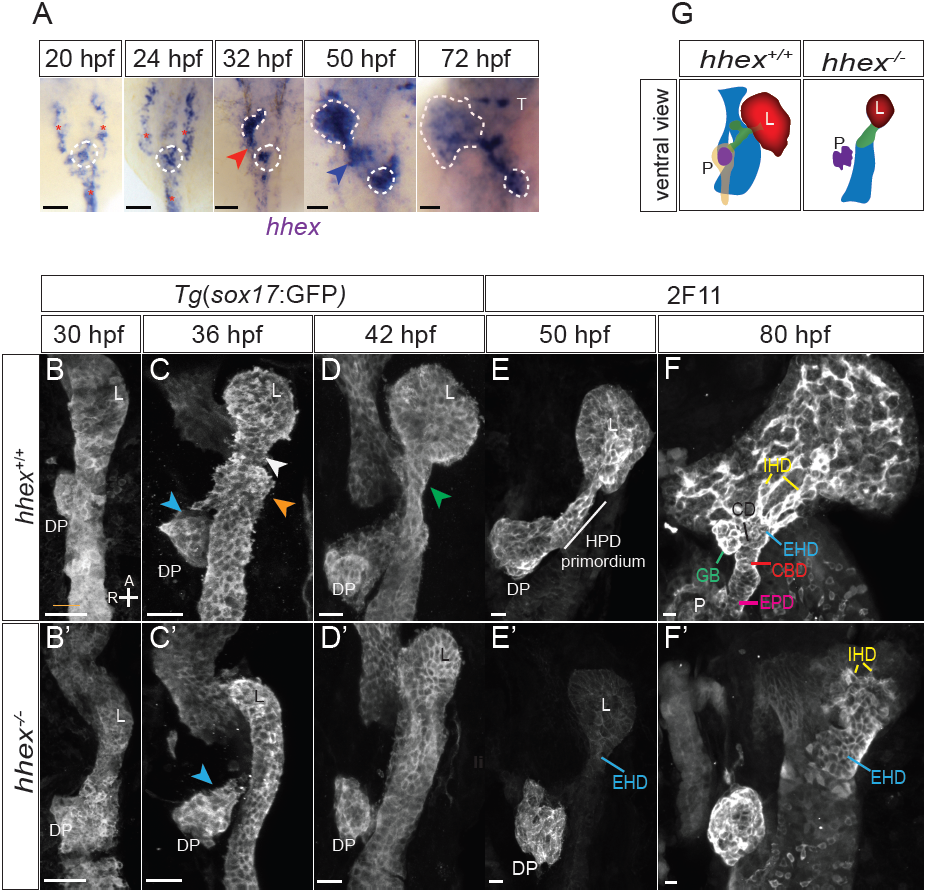
*hhex* regulates the morphogenesis of the HPD system. **(A)** Dorsal views of whole-mount *in situ* hybridization for *hhex* expression. The red and blue arrowheads point to the forming HPD system. The dotted lines highlight the hepatic and pancreatic areas. Red asterisks mark blood endothelial cells. **(B-F’)** Maximum intensity projections of wild-type and *hhex* mutant larvae from 30 to 80 hpf. The HPD system fails to form in *hhex* mutants. Different populations of cells located at three different locations combine to give rise to the HPD system. White arrowhead points to cells caudal to the liver, blue arrowheads point to cells within the dorsal pancreas and orange arrowhead points to cells coming from the anterior pancreatic bud. The green arrowhead highlights the folding of the endoderm into a tube, which extends to form the HPD primordium. All ventral views, anterior to the top.**(G)** Cartoon displays details of the HPD system at 80 hpf in wild-type and *hhex* mutants. The HPD system is in green. A, anterior; CBD, common bile duct; DP, dorsal pancreas; EHD, extra hepatic duct; EPD, extra pancreatic duct; GB, gallbladder; HPD, hepatopancreatic ductal; IPD, intrapancreatic ducts; L, liver; P, pancreas; R, right; T, thymus. Scale bars: 10 μm.

To follow in detail the development of the HPD system, we examined the morphology of the endoderm by analyzing *Tg(sox17*:GFP) expression in the progeny of incrosses of *Tg(sox17:GFP); hhex*^*+/-*^ fish. The *Tg(sox17:GFP)* line labels endodermal cells during the time of early endoderm formation [26, 27] until about 48 hpf when the GFP signal starts to decrease. Therefore, we assessed the morphology of the HPD system at later stages by staining with the 2F11 antibody, which labels the ductal cells [8, 28, 29]. Our confocal analysis provided details of the morphological events that shape the HPD system. At 30 hpf, the liver and dorsal pancreas have budded out (**Fig. 2B**), and by 36 hpf the HPD system starts to form (**Fig. 2C**). Based on their physical location, we can distinguish three populations of cells that will contribute to the formation of the HPD primordium: one located just caudal to the liver (white arrowhead), another one that extends with the anterior pancreatic bud (orange arrowhead), and the third population which is located within the dorsal pancreatic area (blue arrowhead) (**Fig. 2C**). The HPD primordium appears to form in an anterior to posterior fashion, as ductal cells located caudal to the liver start to organize into a tube (**Fig. 2D,** green arrowhead), while cells closer to the dorsal pancreas have not yet done so. By 50 hpf, a single tube, known as the HPD primordium, forms and connects the liver and the pancreas (**Fig. 2E**). During the formation of the HPD primordium, the liver distinctly disassociates from the gut tube and projects ventrally. By 80 hpf, the HPD primordium remodels, and all the constituents of the HPD system become clearly identifiable (**Fig. 2F**).

Our analysis revealed that *hhex* mutants do not develop an HPD system (**Fig. 2B’-G**). At 30hpf the liver and dorsal pancreas budded out in *hhex* mutants as in wild-type siblings, albeit their foregut appears thinner (**Fig. 2B’**). However, by 36 hpf, it became apparent that *hhex* mutants lack a large majority of the cells that give rise to the HPD system (**Fig. 2C’**). There are a few remnant cells, extending either from the liver area or the dorsal pancreatic area, that appear to try to form the HPD system (**Fig. 2C’,** blue arrowhead). However, these cells progressively fail to assemble into an HPD primordium (**Fig. 2D’-F’**). *In vivo* time-lapse confocal imaging of *Tg(gut:GFP)* wild-type and *hhex* mutant embryos show the dynamics of this process (**movie 1** and **movie 2**).

To determine which specific components of the HPD system are dependent on Hhex function, we performed 2F11 immunostaining on 80 hpf larvae from *hhex*^*+/-*^ incrosses. Our analysis revealed that *hhex* mutants lack the gallbladder, CD, CBD, and EPD (**Fig. 2F’**). Nevertheless, a primitive EHD appears to form (**Fig. 2E’** and **2F’**) suggesting that the population of cells located caudal to the liver remained in *hhex* mutants and was able to organize into an EHD. The primitive EHD found in *hhex* mutants was positive for 2F11 as well as for Prox1a, a liver marker that also marks the developing HPD system [8, 28, 30], as detected with the *Tg(prox1a:RFP)* zebrafish line (**Fig. S3**). In wild-type fish at 44 hpf, *prox1a*:RFP^+^ cells are clearly distinguishable in the liver, dorsal pancreas and HPD primordium (**Fig. S3A**). In *hhex* mutants, *prox1a*:RFP^+^ cells can be found in the liver, dorsal pancreas, ectopically in gut cells (**Fig. S3B,** blue arrow), and in cells located where the EHD usually forms (**Fig. S3B,** white arrow). Given that 2F11^+^ cells are also found in this location (**Fig. 2E’**), we hypothesize that these cells are of ductal nature and give rise to the EHD. At later stages, an EHD protruding from the gut becomes evident in *hhex* mutants (**Fig. 2F’**). However, in contrast to wild-type siblings, the EHD fails to extend to the dorsal pancreas or connect with the rest of the HPD system, and instead it directly attaches to the gut tube (**Fig. 2E-G**).

Interestingly, the dorsal pancreas of *hhex* mutants retained high levels of 2F11 expression even after 50 hpf (**Fig. 2E-F’, S4**). 2F11 staining in the dorsal pancreas is known to decrease after the fusion of the dorsal and ventral pancreas [28]. However, 2F11^+^ cells can be observed inside the principal islet of 5 dpf *hhex*^-/-^, while in *hhex*^*+/+*^, 2F11^+^ cells are located outside the principal islet where they organize to form the ductal network (**Fig. S4A-B**). In addition, some *hhex* mutants display amorphous duct-like protrusions from their principal islet (**Fig. S4C-D**). Our results suggest that *hhex* mutants retain ductal cells in the dorsal pancreas. It is not clear, whether this population will contribute to the EPD or the development of the IPDs under normal developmental conditions. Based on these results, we suggest that the HPD system is comprised of two different progenitor populations: one that is Hhex-dependent and is necessary for the development of most of the HPD constituents (the CD, CBD, and EPD), and a second one that is Hhex-independent and gives rise to the EHD.

### *hhex* regulates the specification of HPD progenitors

Since most of the components of the HPD system appear to be absent in *hhex* mutants, we investigated the specification of HPD progenitors. We performed whole-mount *in situ* hybridization for *prox1a* and *sox9b* expression on 36 hpf larvae from *hhex*^*+/-*^ incrosses. These two genes have been previously reported to mark pancreatobiliary cells [7, 28, 31]. In *hhex* mutants, *prox1a* expression remained high in the dorsal pancreas and neural tube, but it was significantly decreased in the forming liver, prospective ventral pancreas, and HPD region (**Fig. 3A-A’,** black arrow). Whole-mount *in situ* hybridization for *sox9b* expression showed that it was unaltered in the dorsal pancreas and neural tube (**Fig. 3B-B’,** white arrow), but it was completely absent from the liver, prospective ventral pancreas, and HPD region in *hhex* mutants (**Fig. 3B-B’,** black arrow). Our results indicate that Hhex function is necessary for the specification of Sox9^+^ pancreaticobiliary progenitors. To assess whether the loss of progenitors was caused by changes in cell death, we performed TUNEL staining in 34 hpf embryos from *Tg(sox17*:*GFP*); *hhex*^*+/-*^ incrosses (**Fig. 3C-D’**). We quantified the number of TUNEL^+^ cells and observed no significant difference between control siblings and *hhex* mutants (1.1± 0.3 in control n=26 embryos total, composed of wild-type n=6 and *hhex*^*+/-*^ n=20 compared to 1.6± 0.6 in *hhex* mutants, n=7; p=0.55) (**Fig. 3E**) indicating that an increase in cell death was not responsible for the loss of HPD progenitors in *hhex* mutants.

**Figure 3.**
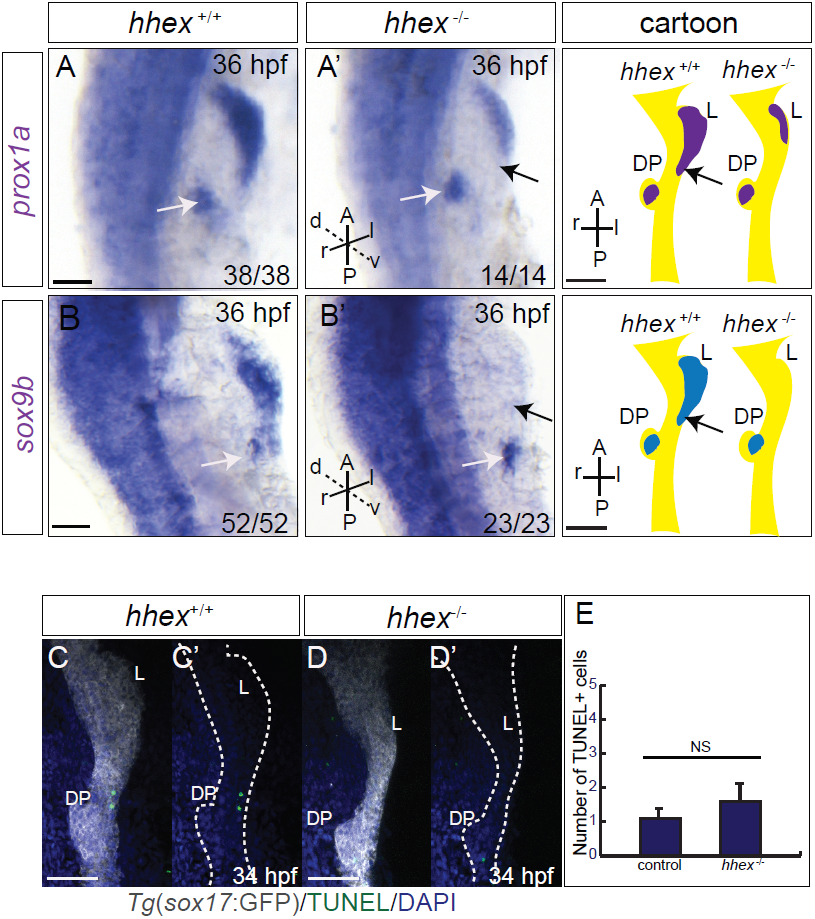
*hhex* regulates the specification of HPD progenitors. **(A-B’)** Dorsal views of whole-mount *in situ* hybridization for **(A-A’)** *prox1a* and **(B-B’)** *sox9b* expression at 36 hpf. *hhex* mutants display reduced *prox1a* and absence of *sox9b* expression in the forming liver, prospective ventral pancreas, and HPD region (black arrows). *prox1a* and *sox9b* expression remained constant in the dorsal pancreas (white arrows) in *hhex* mutants. **(A”**, **B”)** Cartoons of the foregut endoderm display *prox1a* (purple) and *sox9b* (blue) expression in wild-type and *hhex* mutants. Ventral views, anterior to the top. **(C-D’)** Maximum intensity projections of 34 hpf in wild-type sibling and *hhex* mutant stained with TUNEL (green). Endoderm staining is shown in gray and DAPI in blue. Ventral views, anterior to the top. **(E)** Quantification of TUNEL^+^ cells. No significant difference (NS) was observed in the number of TUNEL^+^ cells between control siblings (*hhex*^+/+^, *hhex*^+/-^) and *hhex* mutants. Unpaired t-test was used. Data represented as means ± SEM. A, anterior; d, dorsal; DP, dorsal pancreas; L, liver; r, right; l, left; P, posterior; v, ventral. Scale bars: 50 μm.

### The vasculature is necessary for the patterning of the HPD system, but not for HPD progenitor specification

The dorsal pancreas and liver are specified as early as the ten somite stage [32], while the ventral pancreas is specified between 20 and 24 hpf [33] with the appearance of exocrine marker expression between 29 and 32 hpf [31]. However, the precise developmental window when HPD progenitors are specified remains unknown. Interestingly, *hhex* expression starts in the endoderm around 18-20 hpf, when the first *hhex*^+^ cells appear at the midline and populate the nascent principal islet [34] (**Fig. 2A**). Before that time, *hhex* is strongly expressed in blood endothelial cells during segmentation stages, and in the YSL during gastrulation [35]. Therefore, we questioned whether Hhex regulates HPD progenitor specification cell non-autonomously from any of these tissues.

We first analyzed the contribution of endothelial cells on HPD progenitor specification by assessing the development of the HPD system in *cloche* mutants that lack most endothelial and hematopoietic cells [36, 37]. We reasoned that if Hhex was acting cell non-autonomously from the vasculature, *cloche* mutants should also lack the HPD system. We performed 2F11 immunostaining on *Tg(sox17*:*GFP*); *clo*^*s5/+*^ incrosses and analyzed mutant larvae at 80 hpf. Our results show that although *cloche* mutants developed an HPD system (**Fig. 4**), it is severely dysmorphic and lacks structural distinction between the CBD and the EPD (**Fig. 4B-C**). The EHD is frequently reduced in size and in 43% of the cases the gallbladder is barely distinguishable (**Fig. 4C-C”**). In 57% of *cloche* mutants, there is an increase in the number of ductal cells located in the pancreas. These cells are distributed ectopically around the dorsal pancreas, most commonly locating anterior to the principal islet, where they form amorphous globular structures that disrupt the normal extension of the EPD towards the liver and can block the proper migration of endocrine cells to the principal islet (**Fig. 4B-B”**, data not shown). Our results indicate that blood vessels regulate the patterning of the HPD system and potentially limit its boundaries, but they do not regulate HPD specification.

**Figure 4.**
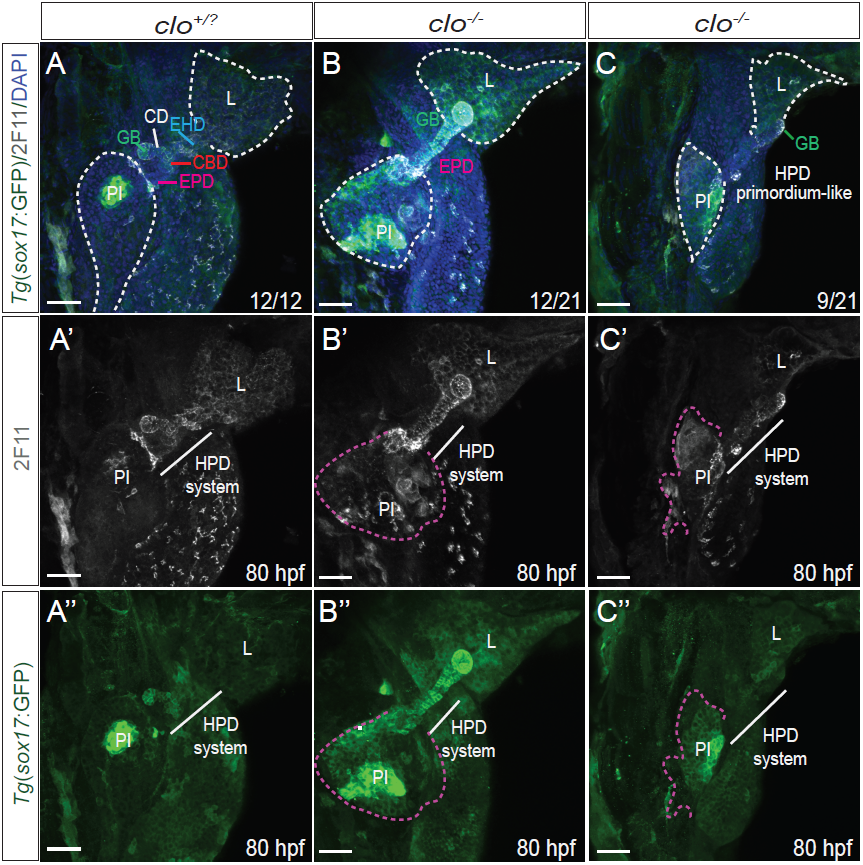
The vasculature is necessary for the patterning of the HPD system, but not for HPD progenitor specification. **(A-C”)** Maximum intensity projections of 80 hpf wild-type siblings and *clo*^*-/-*^ larvae show that the HPD system patterning is dysmorphic in *cloche* mutants. Pink lines highlight the amorphous globular ductal structures present in *cloche* mutants. All ventral views, anterior to the top. CBD, common bile duct; CD, cystic duct; DP, dorsal pancreas; EHD, extra hepatic duct; EPD, extra pancreatic duct; GB, gallbladder; HPD, hepatopancreatic ductal; L, liver; PI, principal islet. Scale bars: 50 μm.

### Signals from the YSL and endoderm regulate the growth of the HPD

*hhex* is highly expressed in the YSL from pre-gastrula stages until the end of gastrulation [35]. The YSL is an important signaling center that has previously been postulated to have a role patterning the anterior pancreatic bud in zebrafish [38]. Therefore, we performed YSL knockdown experiments by injecting *hhex* morpholino [25] at the 1000-cell stage in the YSL (**Fig. 5A-B**). We observed that knockdown of *hhex* in the YSL leads to malformations in the formation of the HPD system with approximately 64% of the larvae at 50 hpf displaying a smaller and thinner HPD system and sporadic cases resembling the phenotype of *hhex* mutants and having a complete absence of the HPD primordium. However, injected embryos frequently display other endodermal defects such as small intestinal bud, liver or laterality defects. In order to test if injections of *hhex* mRNA could rescue the HPD system phenotype, we injected 200pg of *hhex* mRNA to the YSL of 1000-cell stage embryos from *Tg(sox17*:*GFP*); *hhex*^*+/-*^ incrosses. Injections of *hhex* mRNA into the YSL partially rescued the HPD phenotype of *hhex* mutants as assessed by imaging of 50 hpf embryos immunostained for GFP and 2F11 (**Fig. 5C-D**). After *hhex* mRNA injection, bulges of cells were commonly observed caudal to the liver in *hhex* mutants (**Fig. 5D,** blue arrow); and frequently we observed an increase of HPD cells around the principal islet (**Fig. 5D,** white arrow). However, none of the injected *hhex* mutants displayed cells that were incorporated into a fully formed HPD primordium. Our results indicate that reconstituting the function of Hhex in the YSL is not sufficient to direct HPD development and therefore suggest that the function of Hhex in the endoderm is necessary for HPD development.

**Figure 5.**
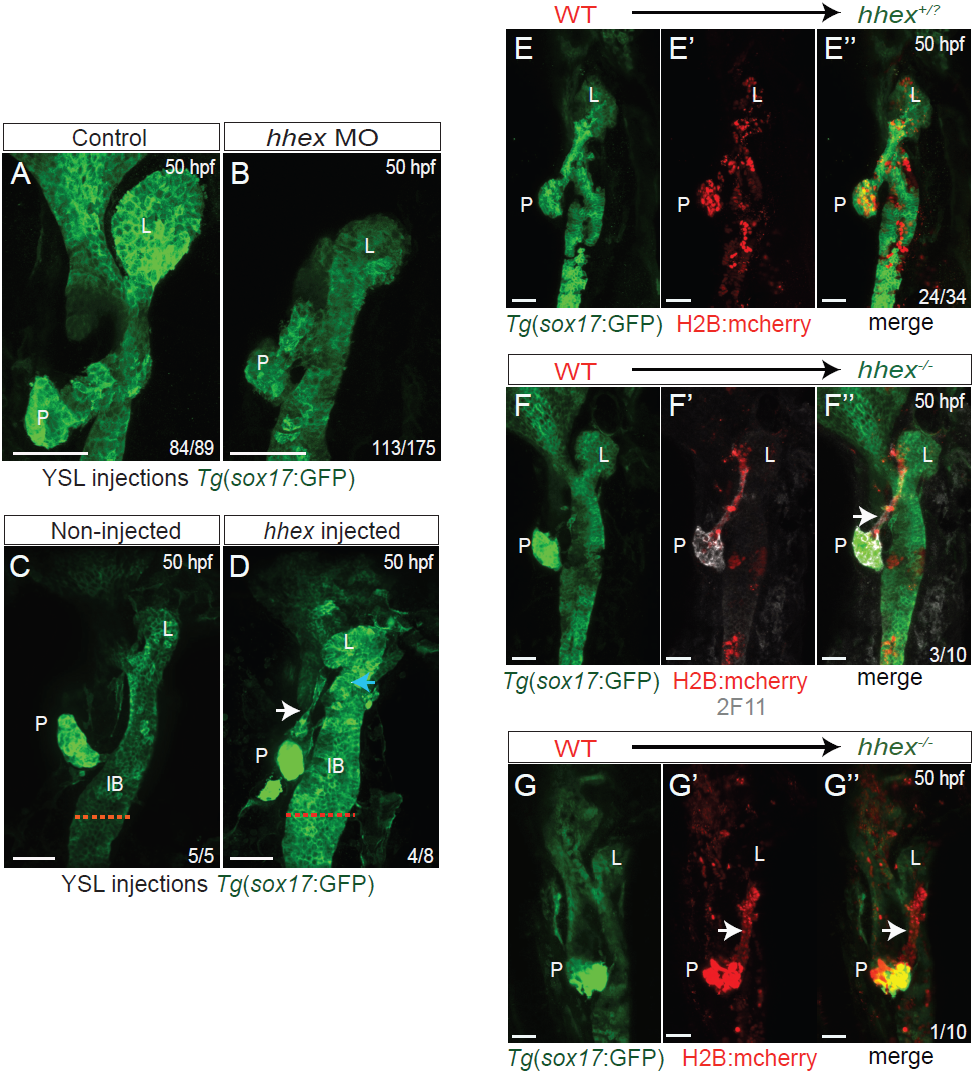
Hhex functions both cell-autonomously and cell non-autonomously to regulate the development of the HPD system. **(A-B)** Maximum intensity projections of 50 hpf control and *hhex* morpholino-injected embryos. Embryos injected with *hhex* MO at the 1000-cell stage resulted in an under-developed HPD system. **(C-D)** Maximum intensity projections of 50 hpf *hhex* mutants non-injected and injected with *hhex* mRNA. Blue arrow points to bulges of cells. White arrow points to additional HPD cells not found in non-injected *hhex* mutants. Red dotted lines highlight the width of the intestinal bud. **(E-G”)** Maximum intensity projections of 50 hpf transplanted wild-type and *hhex* mutant embryos with wild-type donor cells expressing H2B-mcherry. White arrows point to HPD primordium. All ventral views, anterior to the top. IB, Intestinal bulb; L, liver; P, pancreas. Scale bars: 50 μm.

To test the effect of restoring *hhex* function specifically within the endoderm, we performed cell transplantation experiments. We co-injected wild-type donor cells with *sox32* mRNA, to target them to the endodermal fate [39, 40], and with *H2B-mcherry* mRNA to label the transplanted cells. We transplanted cells into host embryos derived from *Tg(sox17*:*GFP*); *hhex*^*+/-*^ incrosses and assessed the incorporation of wild-type cells into the HPD primordium at 50 hpf. Analysis revealed that in wild-type siblings, wild-type donor cells incorporated into the HPD system in 71% of the cases (**Fig. 5E-E”)**. In *hhex* mutants, wild-type donor cells incorporated into the HPD system in 40% of the cases. Strikingly, even though most of the HPD system was absent in mutant hosts, wild-type donor cells were able to position themselves in a diagonal pattern resembling that of the HPD primordium (**Fig. 5F-F”**). In one case, wild-type donor cells organized into a full HPD primordium (**Fig. 5G-G”**), altogether indicating that the endodermal function of Hhex is necessary and sufficient to promote HPD development in a cell-autonomous manner.

## DISCUSSION

Our studies have highlighted a previously unappreciated role for Hhex for instructing multipotent cells of the foregut endoderm into the HPD fate. Here, we show that zebrafish *hhex* mutants are characterized by a complete absence of the HPD primordium and lack further development of the HPD system.While, the CD, CBD, and EPD were unidentifiable in *hhex* mutants, a primitive EHD was still formed. Analysis of s*ox9b*^+^ HPD progenitors showed a complete absence of this cell population in the hepatopancreatic area indicating a role for Hhex in regulating HPD progenitor specification. We propose that the HPD system arises from two populations: one that is responsive to Hhex and necessary for the development of the CD, CBD and EPD and a second population that is Hhex-independent. The contribution of both populations is required for the assembly and development of the mature HPD system.

### Role of the YSL and endoderm in HPD development

During the morphogenesis of the foregut-derived organs in zebrafish, the ventral anterior pancreatic bud fuses with the posterior pancreatic bud by 44 hpf. Cells derived from the anterior bud are known to contribute to the formation of the pancreatic duct, the exocrine tail [41] and probably other ductal components of the HPD system. Before the anterior bud is apparent, the region of the gut endoderm that will give rise to it is located adjacent to the extra-embryonic YSL. Therefore, it has been postulated that the YSL acts as a source of signals for gut endoderm development. Here, we provided evidence of the role of YSL in regulating the growth of the HPD system through the actions of Hhex. Knockdown of *hhex* in the YSL leads to impaired HPD system formation. However, overexpression of *hhex* only partially rescued the *hhex* mutant phenotype, indicating that cell non-autonomous function of Hhex in the YSL is not essential for HPD primordium formation, but it is most likely acting to support the growth of the early foregut.

Our studies show that expression of *hhex* in the zebrafish endoderm can fully rescue the formation of the HPD primordium indicating an essential role for Hhex in the development of the HPD system.Interestingly, in mammals, the specific endodermal deletion of Hhex was reported to lead to the absence of the gallbladder and an abnormal EHD in mammals, with cells of the EHD mostly converting to duodenum-like epithelium [14]. In zebrafish, we observed that the intestinal and ductal boundaries are also disrupted in *hhex* mutants, as intestinal cells appear to express the ductal marker 2F11. The difference in overall HPD system phenotypes between zebrafish and mice may be attributed to differences in species, but alternatively, a disruption of combinatorial function of the extra-embryonic tissue and endoderm might be required for the severity of the phenotype. In mouse, *Hhex* is also expressed in the visceral endoderm in addition to the endoderm raising the question of whether the extra-embryonic function of HHEX might also regulate HPD development in mice. *Hhex* null mice display early embryonic lethality starting from E10.5 [24]. Since the HPD system becomes morphologically distinguishable only after E11.5, the role of HHEX in HPD progenitor specification and development could not be determined. Our data implicates a role for Hhex in the YSL, indicating that signals from both cell non-autonomously from the YSL and cell-autonomous from the endoderm itself contribute to correct specification and formation of the HPD system.

### Role of blood vessels in HPD development

The dynamic cross-talk between the endoderm and endothelial cells orchestrates organ development [42]. For example, paracrine signals from the endothelium are essential for early pancreas differentiation and development [43] and at later stages blood vessels have been reported to negatively regulate pancreatic growth [44]. Our data indicate that signals from the endothelium are not required for HPD progenitor specification; however, we found that they are necessary to pattern the HPD system. Lack of blood vessels produces severely dysmorphic HPD systems with morphologically indistinguishable components that often have an underdeveloped gallbladder. In addition, we found that 30% of *cloche* mutants display ectopic ductal structures in the pancreatic region. Our results suggest that blood vessel-derived signals also negatively regulate the expansion of HPD cells, and hence play a role in establishing HPD system fate boundaries.

The signaling mechanisms by which blood vessels achieve the patterning of the HPD system still need to be determined. Recently, it has been shown that the posterior cardinal vein (PCV) dictates the morphological positioning of the kidney head through the Vegfc/Flt4 pathway [45] and the Vegfc pathway has previously been implicated in dorsal endoderm development and the positioning of the anterior endoderm to the midline [46]. Therefore, the involvement of the Vegfc/Flt-4 signaling in the patterning of the HPD system stands as attractive candidate pathway. Future experiments will help determine the endothelium-derived signaling cues that regulate HPD patterning and morphogenesis.

Our data suggest that Hhex signals from the YSL help support the growth of the HPD system, while Hhex function from the endoderm is essential for the development of the HPD primordium. Based on our data, we thus propose that the HPD develops from two distinct populations of progenitor cells, one that is independent from Hhex function and gives rise to the EHD and likely ductal cells of the pancreas and a second population of progenitors that is dependent on Hhex function and is required for the development of the majority of the components of the HPD system. Finally, our studies also shed light on the important role of blood vessels in HPD patterning and potentially in negatively regulating HPD fate.

## MATERIALS AND METHODS

### Zebrafish mutant and transgenic lines

All zebrafish husbandry was performed under standard conditions in accordance with Institutional (Max Planck Society) and national ethical and animal welfare guidelines. The mutant and transgenic lines used were:

*Tg*(−2.8*fabp10a:EGFP*)^*as*3^[47], abbreviated Tg*(fabp:EGFP*); *Tg(ptf1a:eGFP*) ^*jh*1^ [48];*TgBAC(prox1a:KalT4-4xUAS-E1b:uncTagRFP*)^*nim*5^[49],abbreviated *Tg(prox1a*:*RFP*); *Tg(sox17:GFP*)^*s*870^[27];*Tg(sst2:RFP*)^*gz*19^[50], *Tg(XIEef1a1:GFP*) ^*s854*^ [38], abbreviated *Tg(gut:GFP*); clo^*s*5^ [38]; *hhex*^*s*721^, abbreviated *hhex*^-/-^; and AB fish.

### Generation and genotyping of *hhex* mutants

TALEN arms targeting exon 3 of the *hhex* gene were designed by using the TALEN Targeter 2.0 software by targeting exon 3 of *hhex*. The following TAL effector domains RVDs were used: NI NG HD NG NN NG HD NG HD HD NN HD HD HD NN NI NN and HD NG HD NN HD NG NN NI NN HD NG NN HD NI NN HD NI NG HD NG and assembled using the Golden Gate method [19]. RNA was synthesized with mMESSAGE Machine kit (Ambion), and 100 pg per TALEN arm were injected into one-cell stage zebrafish embryos. Embryos were genotyped by high-resolution melt analysis (HRMA) of PCR products performed in an Eco Real-Time PCR System (Illumina). DyNamo SYBR green (Thermo Fisher Scientific) was used for the PCR and HRMA. The PCR protocol used was 95°C for 10m, 40?cycles of 95°C for 10s, 60°C for 30s, followed by HRMA in the 55-95°C range to distinguished each genotype.

### Immunostaining and Imaging

Embryos were fixed with 4% PFA in Fix Buffer (4% sucrose, 0.15mMCaCl_2_, 0.1M PO_4_ pH7.3) in PBS overnight at 4°C. Immunostaining of whole-mount embryos was performed as previously described [7]. In brief, embryos were deyolked, deskinned, permeabilized with 0.1% Triton for 2hrs, blocked in protein block serum free solution (DAKO) for 4hrs, and incubated overnight with the primary Ab mixture. The concentration for primary antibodies used was as follows: chicken anti-GFP (1:400; Aves), and mouse anti-2F11 (1:50, ABCAM). Secondary antibodies were from Invitrogen and used at a 1:300 concentration. Embryos were washed and incubated with secondary antibody overnight at 4°C. Embryos were then washed and briefly incubated with nuclear stain (DAPI or TOPRO-3). Larvae were mounted on slides and covered with VECTASHIELD antifade mounting medium (vector labs).Alternatively, embryos were mounted in 1% agarose. Samples were imaged using a Zeiss LSM880 with a W Plan Apochromat 20X/NA 0.8DIC.

### Microinjection of morpholinos and Rescue Experiments

For specific knockdown of *hhex* in the YSL: 2.4 ng of *hhex* ATG morpholino GCGCGTGCGGGTGCTGGAATTGCAT [47] and control morpholino CCTCTTACCTCAGTTACAATTTATA from from GeneTools (Philomath, OR) was injected in the YSL at 1000-cell stage in *Tg(sox17*:*GFP*) embryos while for rescue experiments 1000-cell stage embryos from *Tg(sox17*:*GFP*); *hhex*^+/-^ incrosses were injected in the YSL with 200pg of full-length *hhex* mRNA. Larvae were then collected at 50 hpf, stained for GFP, and genotyped by HRMA. Images were captured using a Zeiss LSM880 confocal microscope.

### Transplantations

Donor 1-cell stage AB embryos were co-injected with 150pg of *sox32* mRNA and with 50pg of *H2B-mcherry* mRNA. Host embryos were collected from *Tg(sox17*:*GFP*); *hhex*^+/-^ heterozygous incrosses and 20-50 cells from donors were transplanted in host embryos at mid-blastula stages along the blastoderm margin. Embryos were grown at 28 °C and then they were collected at 50 hpf, stained for GFP and 2F11, and genotyped by HRMA. The contribution of mcherry^+^ cells to the HPD system was determined by confocal microscopy using Zeiss LSM880 confocal microscope.

### TUNEL Assay

Larvae from *Tg(sox17*:*GFP*); *hhex*^-/-^ heterozygous incrosses were fixed in 4%PFA in Fix Buffer (4%sucrose, 0.15mM CaCl_2_, 0.1MPO_4_ pH 7.3) overnight at 4°C. Embryos were washed in PBST and their yolk was manually removed. Endoderm was labeled using anti-GFP and Alexa Fluor 488 staining as described above. Larvae were washed and incubated with 1:300 Alexa Fluor 488 goat anti-chicken (Invitrogen). After the endoderm was labeled, we performed TUNEL Assay using the *in situ* cell death detection kit, TMR red (Roche). Larvae were treated with 0.1% sodium citrate in PBS with 0.3% Triton for 2 min on ice, followed by 1hr incubation at 37°C with TUNEL reaction solution (5 μl TUNEL enzyme and 45 μl TUNEL labeling solution). Samples were then washed three times with PBST, incubated with DAPI, and mounted for imaging. As a positive control, we treated larvae in a DNAse solution (1μl RQ1 DNAse I (Promega), 4 μl DNAse Buffer +35 μl water) for 15 min at 37°C before proceeding with TUNEL labeling reaction. For negative control, we incubated samples with only TUNEL labeling solution. Statistical analyses were performed using unpaired t-test. Embryos were imaged using a Zeiss LSM880 confocal microscope and were collected and genotyped after imaging. Apoptotic cells were quantified using Image J.

## ACKNOWLEDGMENTS

We thank Rita Retzloff and her team for fish care, Sabine Fischer, Nana Fukuda, Sri-Teja Mullapudi and Carmen Büttner for technical and scientific support. We are also grateful to Jenny Pestel for invaluable scientific discussions.

## AUTHOR CONTRIBUTION

A.V., S.G., M.M.C., S.P., and H.H.M. designed and performed experiments. D.Y.R.S. supervised the work. A.V., S.G., M.M.C., and D.Y.R.S. wrote the paper. All authors discussed the results and commented on the paper.

## FIGURE LEGENDS

**Figure S1.**
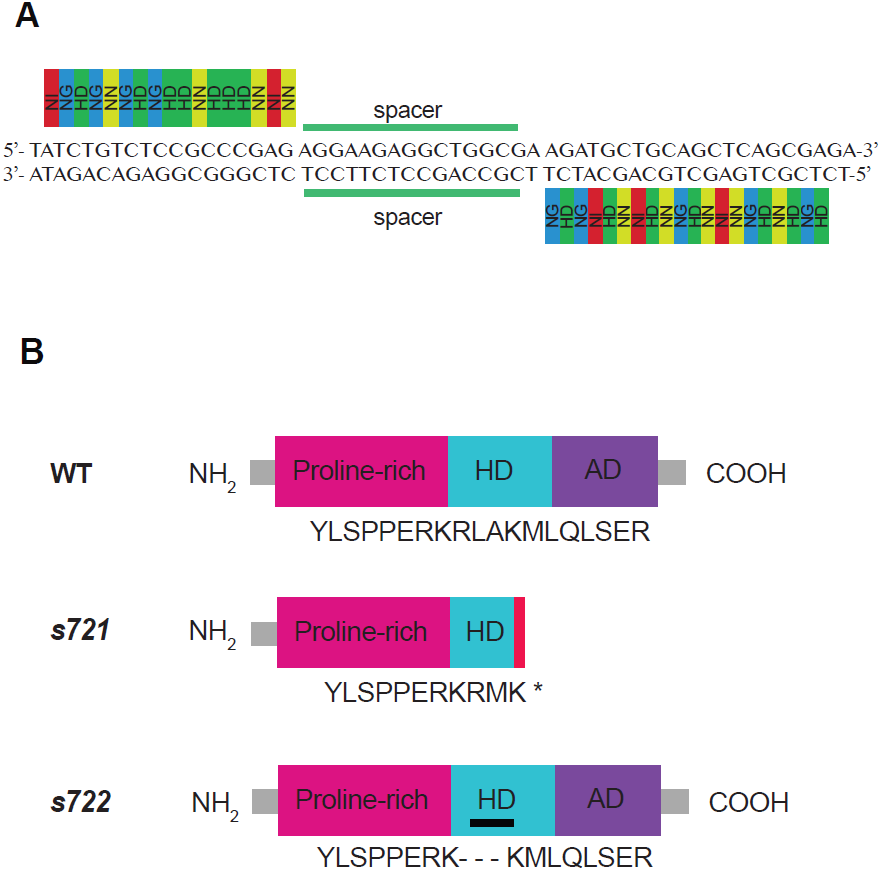
TALEN construct and the different *hhex* alleles. **(A)** TALEN construct for *hhex* mutagenesis. **(B)** Schematic of wild-type and predicted mutant proteins. The *hhex*^*s*721^ lesion leads to a premature stop codon (red bar), and the *hhex*^*s*722^ leads to a three amino acids in-frame deletion within the homeodomain (black bar). AD, acidic domain; HD, homeodomain.

**Figure S2.**
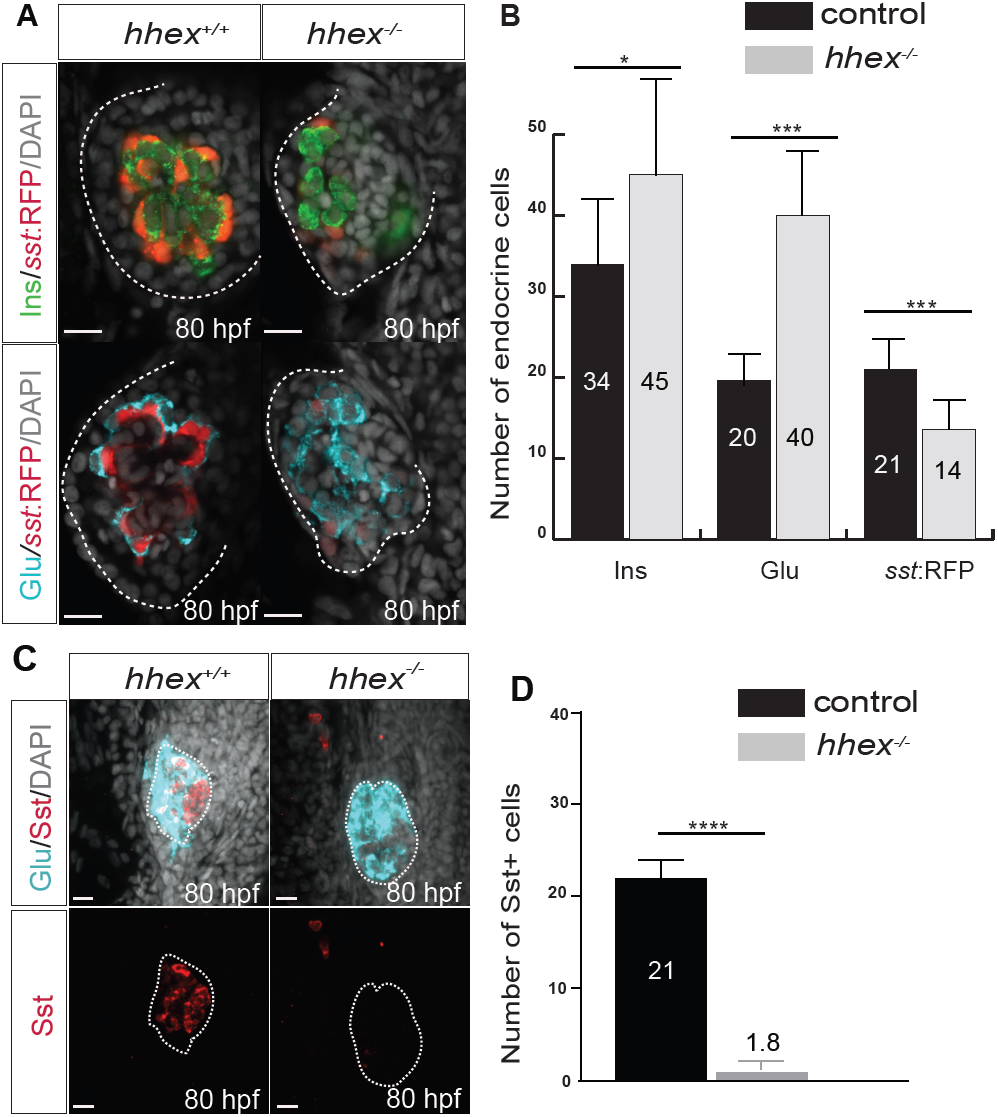
*hhex* mutants display a decrease in the number of Somatostatin^+^ cells. **(A)** Single planes of 80 hpf *hhex*^+/-^ and *hhex*^*-/-*^ principal islet. White borders delineate the pancreas. All lateral views. **(B)** Quantification of the different endocrine cells types: Glucagon (Glu), Insulin (Ins) and s*omatostatin*:RFP^+^ cells (*sst*:RFP). Values are significantly different for all endocrine cells between control siblings (*hhex*^+/+^, *hhex*^+/-^) and *hhex* mutants. (*) p <0.05; (**) p <0.01 and (***) p <0.001. Data represented as means ± SEM. **(C)** Maximum intensity projections of 80 hpf control siblings (*hhex*^+/+^, *hhex*^+/-^) and *hhex* mutant principal islet. White border delineates the principal islet. All lateral views.**(D)** Quantification of Sst^+^ cells immunostained with Sst antibody. Values are significantly different between control siblings and *hhex* mutants. Unpaired t-test was used. Data represented as means ±SEM. Scale bars: 10 μm.

**Figure S3.**
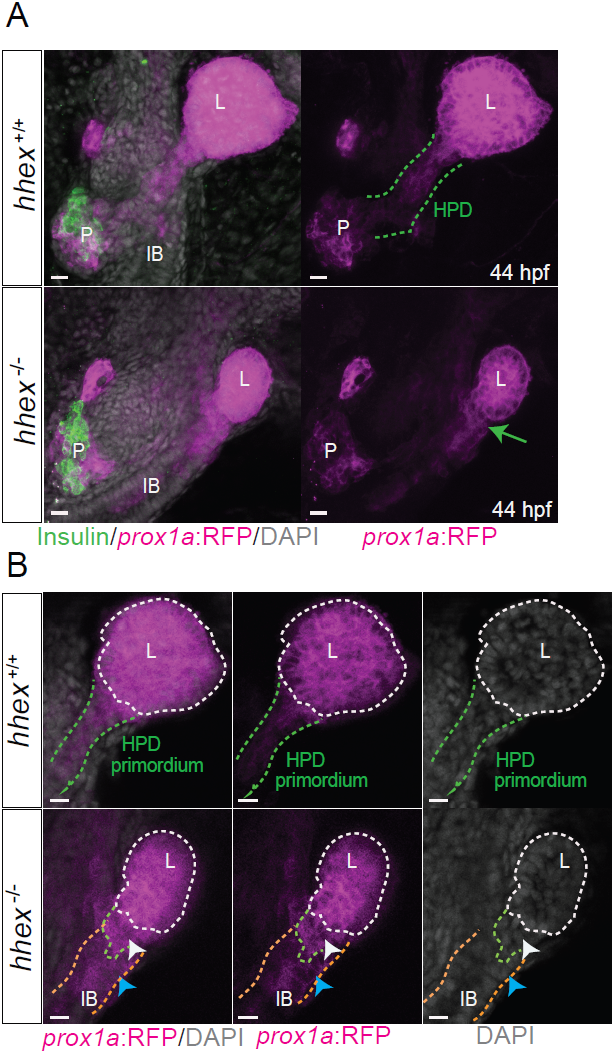
*hhex* mutants develop a primitive extrahepatic duct and display ectopic *prox1a*:RFP expression. **(A)** Maximum intensity projections of 44 hpf *Tg(prox1a:RFP)* embryos stained for Insulin (green) show that *hhex* mutants develop a primitive extrahepatic duct (green arrow). **(B)** Higher magnifications of maximum intensity projections are shown. White arrowheads point to cells located outside the liver boundaries. Blue arrowheads point to gut cells ectopically expressing *prox1a*:RFP in *hhex* mutants. Ventral views, anterior to the top. IB, intestinal bulb; L, liver; P, pancreas. Scale bars: 50μm.

**Figure S4.**
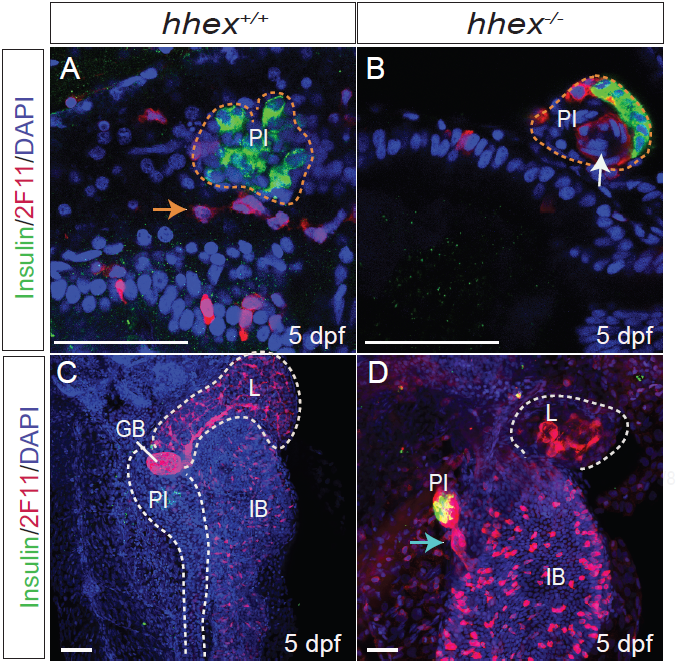
The principal islet of *hhex* mutants retains 2F11^+^ cells. **(A-B)** Single plane images of 5 dpf larvae show that *hhex* mutants display 2F11**^+^** cells inside the principal islet (white arrow), while 2F11**^+^** cells are located outside the islet in *hhex*^+/-^ (orange arrow). Orange dotted lines highlight the principal islet. **(C-D)** Maximum intensity projections of 5 dpf larvae show that some *hhex* mutants display amorphous ductal protusions (blue arrow). White dotted lines highlight the liver and pancreas. Ventral views, anterior to the top. GB, Gallbladder; HPD, Hepatopancreatic ductal; IB, Intestinal bulb; L, liver; PI, principal islet; L, liver. Scale bars: 50 μm.

**Movie 1.** Time-lapse of wild-type *Tg(gut*:*GFP*) zebrafish embryo between 48-55hpf.

**Movie 2.** Time-lapse images of *Tg(gut*:*GFP*); *hhex*^*-/-*^ zebrafish embryo between 48-55 hpf. Note the absence of the ventral pancreatic bud and HPD system.

## REFERENCES

1. Wang, Y., et al., Biliary tree stem cells, precursors to pancreatic committed progenitors: evidence for possible life-long pancreatic organogenesis. Stem Cells, 2013. 31(9): p. 1966–79.

2. Cardinale, V., et al., The biliary tree--a reservoir of multipotent stem cells. Nat Rev Gastroenterol Hepatol, 2012. 9(4): p. 231–40.

3. Carpino, G., et al., Biliary tree stem/progenitor cells in glands of extrahepatic and intraheptic bile ducts: an anatomical in situ study yielding evidence of maturational lineages. J Anat, 2012. 220(2): p. 186–99.

4. Cardinale, V., et al., Multipotent stem/progenitor cells in human biliary tree give rise to hepatocytes, cholangiocytes, and pancreatic islets. Hepatology, 2011. 54(6): p. 2159–72.

5. Semeraro, R., et al., Multipotent stem/progenitor cells in the human foetal biliary tree. J Hepatol, 2012. 57(5): p. 987–94.

6. Strobel, O., et al., Pancreatic duct glands are distinct ductal compartments that react to chronic injury and mediate Shh-induced metaplasia. Gastroenterology, 2010. 138(3): p. 1166–77.

7. Delous, M., et al., Sox9b is a key regulator of pancreaticobiliary ductal system development. PLoS Genet, 2012. 8(6): p. e1002754.

8. Dong, P.D., et al., Fgf10 regulates hepatopancreatic ductal system patterning and differentiation. Nat Genet, 2007. 39(3): p. 397–402.

9. Villasenor, A. and D.Y.R. Stainier, On the development of the hepatopancreatic ductal system. Semin Cell Dev Biol, 2017. 66: p. 69–80.

10. Kanai-Azuma, M., et al., Depletion of definitive gut endoderm in Sox17-null mutant mice. Development, 2002. 129(10): p. 2367–79.

11. Spence, J.R., et al., Sox17 regulates organ lineage segregation of ventral foregut progenitor cells. Dev Cell, 2009. 17(1): p. 62–74.

12. Bort, R., et al., Hex homeobox gene-dependent tissue positioning is required for organogenesis of the ventral pancreas. Development, 2004. 131(4): p. 797– 806.

13. Bort, R., et al., Hex homeobox gene controls the transition of the endoderm to a pseudostratified, cell emergent epithelium for liver bud development. Dev Biol, 2006. 290(1): p. 44–56.

14. Hunter, M.P., et al., The homeobox gene Hhex is essential for proper hepatoblast differentiation and bile duct morphogenesis. Dev Biol, 2007. 308(2): p. 355–67.

15. Bogue, C.W., et al., Hex expression suggests a role in the development and function of organs derived from foregut endoderm. Dev Dyn, 2000. 219(1): p. 84–9.

16. Pelster, B. and W.W. Burggren, Disruption of hemoglobin oxygen transport does not impact oxygen-dependent physiological processes in developing embryos of zebra fish (Danio rerio). Circ Res, 1996. 79(2): p. 358–62.

17. Gut, P., et al., Little Fish, Big Data: Zebrafish as a Model for Cardiovascular and Metabolic Disease. Physiol Rev, 2017. 97(3): p. 889–938.

18. Bedell, V.M., et al., In vivo genome editing using a high-efficiency TALEN system. Nature, 2012. 491(7422): p. 114–8.

19. Cermak, T., et al., Efficient design and assembly of custom TALEN and other TAL effector-based constructs for DNA targeting. Nucleic Acids Res, 2011. 39(12): p. e82.

20. Dahlem, T.J., et al., Simple methods for generating and detecting locus-specific mutations induced with TALENs in the zebrafish genome. PLoS Genet, 2012. 8(8): p. e1002861.

21. Martinez Barbera, J.P., et al., The homeobox gene Hex is required in definitive endodermal tissues for normal forebrain, liver and thyroid formation. Development, 2000. 127(11): p. 2433–45.

22. Zhang, J., et al., The diabetes gene Hhex maintains delta-cell differentiation and islet function. Genes Dev, 2014. 28(8): p. 829–34.

23. Zhao, H., et al., Homeoprotein hhex-induced conversion of intestinal to ventral pancreatic precursors results in the formation of giant pancreata in Xenopus embryos. Proc Natl Acad Sci U S A, 2012. 109(22): p. 8594–9.

24. Keng, V.W., et al., Homeobox gene Hex is essential for onset of mouse embryonic liver development and differentiation of the monocyte lineage. Biochem Biophys Res Commun, 2000. 276(3): p. 1155–61.

25. Wallace, K.N., et al., Zebrafish hhex regulates liver development and digestive organ chirality. Genesis, 2001. 30(3): p. 141–3.

26. Chung, W.S. and D.Y. Stainier, Intra-endodermal interactions are required for pancreatic beta cell induction. Dev Cell, 2008. 14(4): p. 582–93.

27. Alexander, J. and D.Y. Stainier, A molecular pathway leading to endoderm formation in zebrafish. Curr Biol, 1999. 9(20): p. 1147–57.

28. Zhang, D., et al., Identification of Annexin A4 as a hepatopancreas factor involved in liver cell survival. Dev Biol, 2014. 395(1): p. 96–110.

29. Crosnier, C., et al., Delta-Notch signalling controls commitment to a secretory fate in the zebrafish intestine. Development, 2005. 132(5): p. 1093–104.

30. Noel, E.S., et al., Organ-specific requirements for Hdac1 in liver and pancreas formation. Dev Biol, 2008. 322(2): p. 237–50.

31. Manfroid, I., et al., Zebrafish sox9b is crucial for hepatopancreatic duct development and pancreatic endocrine cell regeneration. Dev Biol, 2012. 366(2): p. 268–78.

32. Chung, W.S., C.H. Shin, and D.Y. Stainier, Bmp2 signaling regulates the hepatic versus pancreatic fate decision. Dev Cell, 2008. 15(5): p. 738–48.

33. Naye, F., et al., Essential roles of zebrafish bmp2a, fgf10, and fgf24 in the specification of the ventral pancreas. Mol Biol Cell, 2012. 23(5): p. 945–54.

34. Wallace, K.N. and M. Pack, Unique and conserved aspects of gut development in zebrafish. Dev Biol, 2003. 255(1): p. 12–29.

35. Ho, C.Y., et al., A role for the extraembryonic yolk syncytial layer in patterning the zebrafish embryo suggested by properties of the hex gene. Curr Biol, 1999. 9(19): p. 1131–4.

36. Reischauer, S., et al., Cloche is a bHLH-PAS transcription factor that drives haemato-vascular specification. Nature, 2016. 535(7611): p. 294–8.

37. Stainier, D.Y., et al., Cloche, an early acting zebrafish gene, is required by both the endothelial and hematopoietic lineages. Development, 1995. 121(10): p. 3141–50.

38. Field, H.A., et al., Formation of the digestive system in zebrafish. I. Liver morphogenesis. Dev Biol, 2003. 253(2): p. 279–90.

39. Kikuchi, Y., et al., casanova encodes a novel Sox-related protein necessary and sufficient for early endoderm formation in zebrafish. Genes Dev, 2001. 15(12): p. 1493–505.

40. Aoki, T.O., et al., Molecular integration of casanova in the Nodal signalling pathway controlling endoderm formation. Development, 2002. 129(2): p. 275–86.

41. Field, H.A., et al., Formation of the digestive system in zebrafish. II. Pancreas morphogenesis. Dev Biol, 2003. 261(1): p. 197–208.

42. Villasenor, A. and O. Cleaver, Crosstalk between the developing pancreas and its blood vessels: an evolving dialog. Semin Cell Dev Biol, 2012. 23(6): p. 685– 92.

43. Yoshitomi, H. and K.S. Zaret, Endothelial cell interactions initiate dorsal pancreas development by selectively inducing the transcription factor Ptf1a. Development, 2004. 131(4): p. 807–17.

44. Magenheim, J., et al., Blood vessels restrain pancreas branching, differentiation and growth. Development, 2011. 138(21): p. 4743–52.

45. Chou, C.W., et al., The endoderm indirectly influences morphogenetic movements of the zebrafish head kidney through the posterior cardinal vein and VegfC. Sci Rep, 2016. 6: p. 30677.

46. Ober, E.A., et al., Vegfc is required for vascular development and endoderm morphogenesis in zebrafish. EMBO Rep, 2004. 5(1): p. 78–84.

47. Her, G.M., et al., In vivo studies of liver-type fatty acid binding protein (L-FABP) gene expression in liver of transgenic zebrafish (Danio rerio). FEBS Lett, 2003. 538(1-3): p. 125–33.

48. Godinho, L., et al., Targeting of amacrine cell neurites to appropriate synaptic laminae in the developing zebrafish retina. Development, 2005. 132(22): p. 5069–79.

49. van Impel, A., et al., Divergence of zebrafish and mouse lymphatic cell fate specification pathways. Development, 2014. 141(6): p. 1228–38.

50. Li, Z., et al., Generation of living color transgenic zebrafish to trace somatostatin-expressing cells and endocrine pancreas organization. Differentiation, 2009. 77(2): p. 128–34.

